# Continuous monitoring of plant transpiration dynamics with a leaf-mounted sensor across environmental conditions

**DOI:** 10.64898/2026.06.15.732311

**Authors:** Marcela T. Miranda, Luciano Pereira, Swetlana Kreinert, Jonas Ott, Antônia F. D. Fernandes, Camila C. Carvalho, Steven Jansen, Rafael V. Ribeiro

## Abstract

- Transpiration plays a central role in plant water relations and strongly influences plant growth. Continuous monitoring is essential for understanding responses to environmental conditions and improving water management in both natural and agricultural systems. Gas-exchange techniques such as infrared gas analysers (IRGAs) and porometers are widely used but are challenging for long-term or large-scale monitoring. On the other hand, the FylloClip is a low-cost, leaf-mounted capacitance sensor developed previously to monitor transpiration by detecting condensation of water vapour near the leaf surface. Here, we evaluated the potential of the FylloClip for monitoring transpiration dynamics and assessed environmental conditions that may affect its performance.
- The FylloClip was tested under growth chamber, greenhouse, and tropical field conditions. We evaluated how its capacitance measurements respond to rainfall, temperature and humidity, and compared FylloClip measurements with transpiration measured with an IRGA.
- There was a strong correlation (*r* = 0.85) between FylloClip and IRGA data. Both systems captured similar diurnal transpiration patterns, with transpiration declining simultaneously under water deficit. Rainfall and very high relative humidity produced FylloClip signals that could be misinterpreted as high transpiration, although transpiration is negligible under these conditions.
- Our results revealed that FylloClips capture temporal patterns of transpiration with high accuracy and resolution, providing a reliable tool for long-term, large-scale monitoring of transpiration dynamics in ecophysiological studies and precision agriculture.

## Introduction

When stomata in leaves are open for CO_2_ uptake during photosynthesis, water vapour from the moist leaf mesophyll is lost to the atmosphere in a process called transpiration. Transpiration is not risky for plants under well-hydrated conditions. Yet leaf transpiration may lead to an imbalance in the plant water status, which can affect plant development, and lead to tissue desiccation and damage of the photosynthetic apparatus (Fortunel *et al*., 2023). In fact, drought is a major yield- and growth-limiting factor for plants, especially considering climate change (Dietz *et al*., 2021).

Agriculture, particularly crop production, represents the largest human use of freshwater, primarily due to irrigation, which drives plant transpiration and evaporation from soils (Campbell *et al*., 2017). Climate change driven by human activities is altering temperature and precipitation patterns across many regions worldwide at an unprecedented rate. As a consequence, extreme climate events, including heat waves, droughts, and storms, are becoming more frequent and severe, with significant implications for natural and agricultural systems (Yang *et al*., 2024). Besides the influence of soil water availability, transpiration can also be modulated by several environmental factors, including air temperature, photosynthetically active radiation (PAR), relative humidity, and wind (Drake *et al*., 1970; Zhu *et al*., 2022). In this context, accurate monitoring of plant transpiration at high spatial and temporal resolution, and across both individual plants and ecosystem scales, is essential for understanding plant responses to environmental factors.

High-precision sensors have become increasingly important in modern agriculture and forest management, enabling real-time monitoring and continuous data collection (Ruan *et al*., 2019; Chen *et al*., 2023). Ground-based sensors and wireless sensor networks allow the monitoring of environmental conditions and dynamic plant physiological responses, improving management decisions and a more efficient use of natural resources (López Riquelme *et al*., 2009; Zhang *et al*., 2023; Chaplin *et al*., 2025). Beyond their application in agriculture and forestry, collecting large, sensor-based datasets opens exciting and novel opportunities for scientists, allowing robust, comprehensive exploration of a wide range of plant processes at unprecedented resolution (Pereira *et al*., 2020; Gleason *et al*., 2024). In addition, data with a high temporal resolution allow the identification of subtle patterns and correlations that might go unnoticed in coarser-resolution datasets (Miranda *et al*., 2024).

Gas-exchange analysers became widely used in plant ecophysiology since the 1980s, especially after the introduction of portable models that allowed for field measurements (Long *et al*., 1996). Based on the tight relationship between leaf CO_2_ assimilation and water losses by transpiration, leaf gas exchange can be monitored using infrared gas exchange analysers (IRGAs). As IRGAs monitor changes in water vapour and CO_2_, the estimation of net photosynthesis and transpiration *in vivo* is possible (Douthe *et al*., 2018). Although the IRGAs are important tools for measuring plant gas exchange, their applications in continuous monitoring are limited due to high costs, the need for specialized operation, and the logistical challenges of deploying such devices across multiple sites or over extended periods. These constraints reduce their practicality for large-scale or long-term studies, particularly under field conditions.

The FylloClip is a low-cost, leaf-mounted capacitance sensor developed and tested by Thalheimer (2023) to monitor changes in plant water dynamics at the leaf level. Basically, the device consists of a printed circuit board (PCB) with a circular capacitive sensor, measuring variations in sensor capacitance when water accumulates by condensation on the sensor surface. At high transpiration rates, a high amount of water vapour wets the sensor surface, resulting in high capacitance readings. On the contrary, when stomata close, the amount of vapour available for condensation is low, leading to low sensor capacitance. However, the FylloClip signal described by Thalheimer *et al*. (2026) was suggested not to represent transpiration alone, but rather reflects the dynamic balance between condensation and evaporation processes occurring at the leaf–sensor interface.

To our knowledge, the performance of the FylloClip sensor has not yet been evaluated under a wide range of environmental conditions, and no direct comparison has yet been made between FylloClip capacitance measurements and leaf transpiration based on traditional gas exchange methods. In this context, the aim of this study was to assess the potential of the FylloClip as a tool for monitoring plant transpiration, and to identify environmental conditions that may limit its universal application. To achieve this, FylloClip measurements were compared with transpiration data taken with an IRGA, which served as a benchmark for interpreting the sensor signals. The sensor was then tested under a range of controlled and field conditions. These experiments allowed us to explore how variations in environmental parameters affect sensor responses and to identify potential limitations.

## Material and Methods

### FylloClip construction and measurements

The FylloClips were constructed according to Thalheimer (2023). Details and instructions are available on the FylloClip GitHub repository (https://github.com/FylloClip; for additional information, please contact the authors). The device consists of a printed circuit board with a circular interdigitated capacitive sensor plate with a 2 cm diameter (Fig. 1). Water condensation on sensor surface modifies the dielectric properties of the medium surrounding the electrodes, changing the FylloClip signals. We made small adaptations for data transmission using an ESP8266 microcontroller to transmit the sensor data to a computer instead of using a LoRaWAN module. In addition, we used a BME280 sensor (Bosch, Germany) to monitor air relative humidity and temperature, which we used to calculate the dew point temperature and air vapour pressure deficit (VPD). In all experiments, one FylloClip sensor was installed per plant on a fully expanded, mature, and non-shaded leaf. Once installed, the FylloClip sensors were not repositioned or reinstalled and remained attached to the same leaf throughout the experimental period. The FylloClip measurements were taken every 15 minutes, and the capacitance measurements were transformed into relative values, with 100% being the maximum value measured. The relative capacitance (RC, %) measured with the FylloClip was estimated as:

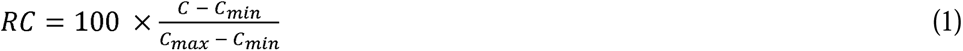

**Figure 1:**
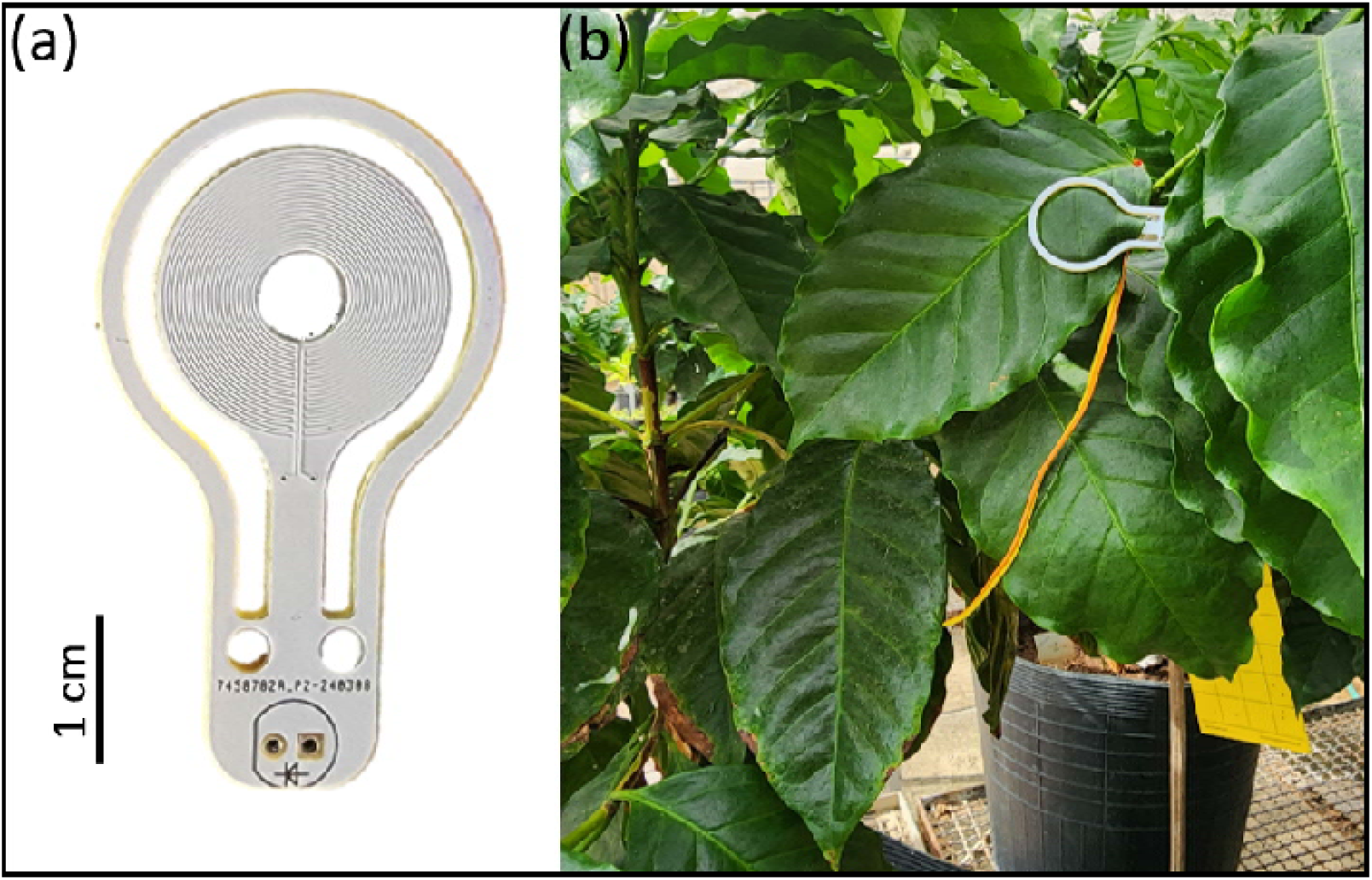
(a) a FylloClip sensor showing the interdigitated electrode geometry on a printed circuit board; the scale bar represents 1 cm. (b) a FylloClip sensor installed on a coffee plant, with the sensor clipped to the abaxial leaf surface.

where *C* is the capacitance measured with the FylloClip, *C*_min_ and *C*_max_ are the minimum and maximum capacitance measurements taken with the FylloClip during the entire experimental period.

This normalisation procedure was necessary because absolute capacitance values vary among individual sensors due to device-specific offsets and baseline differences, most likely due to differences in sensor manufacturing, which prevent direct comparison of raw measurements. By normalising the data of each sensor, FylloClip measurements became directly comparable across sensors, while preserving the temporal dynamics of the signal.

### Experiment 1: FylloClip measurements in maize plants inside a growth chamber

#### Plant material and experimental design

Plants of maize (*Zea mays* L. cv. ‘Damaun KS’) were cultivated under greenhouse conditions in four litre pots filled with a commercial substrate, with daily irrigation. Ten days before the experiment started, plants were transferred to an LED-equipped growth chamber (Polyklima, Germany), with controlled air humidity, temperature, and PAR to minimise environmental variation. The chamber was programmed for a 12-hour photoperiod (06:00–18:00 h), with a 45-minute transition between light and dark to simulate sunrise and sunset. The environmental setpoints were 23 °C and RH 65%, with PAR of 600 μmol m□² s□¹. The FylloClips were installed on eight plants when they were 35 days old, with one sensor per plant. Measurements were taken every 15 minutes for eight days, and the plants were irrigated daily. Although a higher temporal resolution with FylloClips would be feasible, the 15 min intervals were adequate for our comparison with IRGA-data.

### Experiment 2: Comparison of FylloClip and IRGA measurements taken in rice, common bean, and coffee seedlings

#### Plant material and experimental design

This experiment was conducted inside a room with natural fluctuation of air temperature and RH, i.e., no air-conditioning. The plants were put under an LED light source programmed for a 12-hour photoperiod (6:00–18:00h) with PAR at ∼480 μmol m^−2^ s^−1^. Plants of rice (*Oryza sativa* L. cv ‘BRS Pampeira’), common bean (*Phaseolus vulgaris* cv. ‘IAC 1849 Polaco’), and coffee (*Coffea arabica* cv. ‘Catuai Vermelho’) were cultivated in three litter pots filled with commercial substrate. Plants remained under greenhouse and daily irrigation, before moving them to the room conditions. As we had only one IRGA device for comparisons with FylloClip data, species were evaluated one at a time. Rice experiments were conducted in March, common bean in May, and coffee in August 2025. Due to these temporal differences, environmental conditions inside the room (temperature and RH) varied when comparing the room conditions for each species (Supplementary Material). One week before starting the measurements, plants were placed under the light source for acclimation, and remained there throughout the measurements. When the measurements started, rice plants were 10 weeks old, common bean plants were 22 days old, and coffee plants were six months old. Coffee plants were maintained well irrigated during the experimental period (eight days). In contrast, rice and common bean plants were well irrigated for six days, after which irrigation was suspended until transpiration reached values close to zero. This period lasted three days for rice, and five days for common bean.

#### Leaf gas exchange

Transpiration was measured every 15 minutes with an infrared gas analyser (LI□6400XT, LI□COR, USA) under ambient CO_2_ concentration. The IRGA chamber was configured to measure the ambient light intensity, and automatically adjusted its internal light source to match this intensity, thereby ensuring that the leaf inside the chamber was at the same light level as it would be outside the chamber. The IRGA and the FylloClip were clamped on the same leaf, with the IRGA closer to the basal part of the leaf. For a correlation analysis between transpiration and capacitance measurements, transpiration values were converted to relative values (using the same minimum and maximum rescaling approach described before) to ensure comparability between variables expressed on different absolute scales.

### Experiment 3: Coffee plants under greenhouse conditions

#### Plant material and experimental design

For this experiment, we used *Coffea arabica* cv. ‘Catuai SH3’ planted in 25 L pots filled with commercial substrate. Plants were maintained under greenhouse conditions without temperature and RH control, and under natural light fluctuation. This experiment was designed to evaluate the FylloClip under conditions of larger environmental variation than the settings of Experiment 2. The plants were one year old and were irrigated daily during the experimental period. One FylloClip was installed per plant on a fully expanded, sun-exposed leaf, with a total of eight plants monitored. Data were collected every 15 minutes for 15 days from March 15 to 29, 2025.

### Experiment 4: Coffee plants under field conditions

#### Plant material and experimental design

*Coffea arabica* cv. ‘Catuai Vermelho’ plants were cultivated in 10 L pots filled with commercial substrate and grown outdoors at the State University of Campinas, Campinas, SP, Brazil (22°82’ S, 47°07’ W, 599 m a.s.l.). The plants were one year old and one FylloClip per plant was installed on a fully expanded, sun-exposed leaf. Since our goal was to monitor transpiration under field conditions, and not to investigate intraspecific variation, we monitored four plants. Data were collected every 15 minutes for 30 days from December 19, 2025 to January 17, 2026. During the experimental period, plants were under rainfed conditions. Environmental data (air temperature, RH, and rainfall) were obtained from a meteorological station located within the campus and provided by the Center for Meteorological and Climatic Research Applied to Agriculture (CEPAGRI/UNICAMP).

### Data analysis

In all experiments, we applied a three-point moving average corresponding to a 45-minute window, based on the average value of three consecutive FylloClip measurements. Therefore, each 15-minute data point was averaged as the combined value of this actual measurement, the previous, and the following one. This approach was done to reduce high-frequency sensor noise and to improve the visualisation of temporal patterns in graphs. However, this smoothing procedure was not used for any statistical analyses or model fitting.

In experiments 1, 3 and 4, mean relative capacitance values were calculated, and standard errors were computed accordingly. For the correlation analysis in experiment 2, IRGA and FylloClip data were paired according to the closest measurement time between instruments. To avoid artificial inflation of the correlation between capacitance and transpiration due to the high number of near-zero values recorded at night, we performed the analysis using both the full dataset and a subset that was restricted to daytime data only. At night, both measurements tend to approach zero, which could bias the correlation by overrepresenting low-variation points. Therefore, the daytime analysis was used to assess whether the IRGA-FylloClip agreement was robust.

The dew point temperature was calculated using the {weathermetrics} package in R, and the difference between air temperature and dew point temperature (ΔT, hereafter referred to as dew point depression) was computed. Air vapour pressure deficit (VPD) was calculated using the {plantecophys} package. For comparative purposes, the ΔT scale was fixed between 0 and 18 °C for all graphs, covering the full ΔT range observed across all experiments. We chose to present the data and the corresponding ΔT values because of the direct relationship of ΔT with condensation, which would affect the FylloClip measurements. VPD data are provided in the Supplementary Material (Figs. S1 to S4), as biological driver behind transpiration. The relationship between relative capacitance (RC) and ΔT was evaluated using segmented regression to identify potential changes in the response of the sensor signal to ΔT. Only night-time data were included in such analysis, as transpiration was assumed to be minimal during this period, and variation in RC was therefore attributed primarily to ΔT effects. A linear model was first fitted between RC and ΔT, and a breakpoint was then estimated using the {segmented} package in R. The segmented regression fits two linear relationships separated by an estimated breakpoint, which is calculated by minimizing the residual sum of squares. The estimated breakpoint represented the ΔT value at which the slope of the RC–ΔT relationship changes. All data were analysed using R (R Core Team, 2025).

## Results

In the first experiment with maize plants under controlled conditions, the FylloClips presented similar patterns among the eight plants (Fig. 2). Shortly after the lights turned on inside the growth chamber, there was an increase in the sensor capacitance that remained stable during the day. When the light was switched off, there was a decrease in the FylloClip records. During days 3, 5, 6, and 7 of the experiment, temporary reductions in RC were observed (Fig. 2). These fluctuations were likely caused by unintended disturbances to the plants during irrigation and/or by the repeated opening and closing of the chamber, which may have affected the sensor–leaf contact and shortly altered the capacitance readings. During the experiment, the air temperature ranged from 22.5 to 23.5 °C and RH from 62 to 67% (Fig. S1). Since air RH and temperature were controlled inside the chamber, the dew point temperature did not change much, with ΔT varying from 6.4 to 7.8 °C and VPD ranging from 0.9 to 1.1 kPa during the experimental period. No guttation was observed on maize leaves in the early morning.

**Figure 2:**
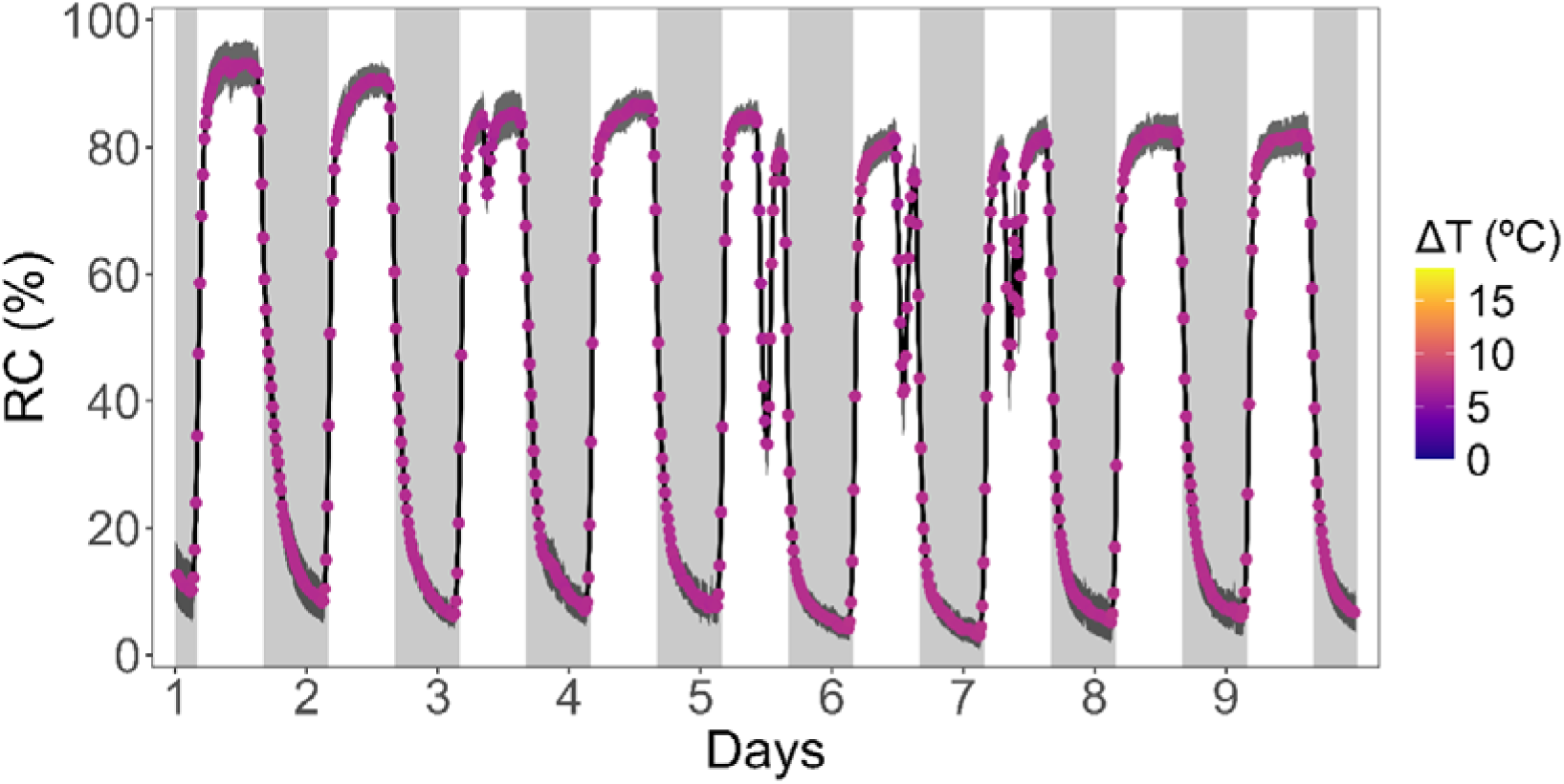
Relative capacitance (%) monitored with a FylloClip every 15 minutes in maize plants under controlled conditions in a growth chamber over nine consecutive day-night cycles. Each measurement represents the mean values (*n* = 8 plants, with 1 sensor per plant) and was smoothed using a three-point moving average. The dark grey shadow represents the standard error. Symbols are coloured according to the dew point depression value (ΔT), and the pink colour shows that there was no daily variation in ΔT. Grey background areas represent dark periods (night), while white areas represent periods when the light was on (day).

Experiment 2 was conducted separately for each species due to the availability of one single IRGA. Environmental conditions, therefore, varied among the experiments. For rice plants, room temperature varied from 26.3 to 32.3 °C and RH from 43 to 70% (Fig S2a), resulting in ΔT values between 5.8 and 14 °C and VPD varying from 2 to 4 kPa (Fig S2b). For common bean plants, room temperature ranged from 19.4 to 26.3 °C, and RH from 46 to 76% (Fig S2c), with ΔT varying from 4.3 to 12 °C and VPD ranging from 1.3 to 2.5 kPa (Fig S2d). Finally, room temperature ranged from 18 to 23 °C and RH from 39 to 68% for coffee plants (Fig S2e), corresponding to ΔT values between 6.2 and 14.2 °C and VPD values between 1.8 and 3 (Fig S2f). For the three species studied, the daily pattern of capacitance measured with the FylloClip followed a similar pattern to that of transpiration measured with the IRGA. When the light was turned on, there was a rapid increase in leaf transpiration and sensor capacitance, followed by a decrease when the light was turned off (Fig. 3). The maximum value of leaf transpiration measured with the IRGA in rice plants was 5.6 mmol m^−2^ s^−1^, while for common bean and coffee plants was 3.6 and 0.93 mmol m^−2^ s^−1^, respectively.

**Figure 3:**
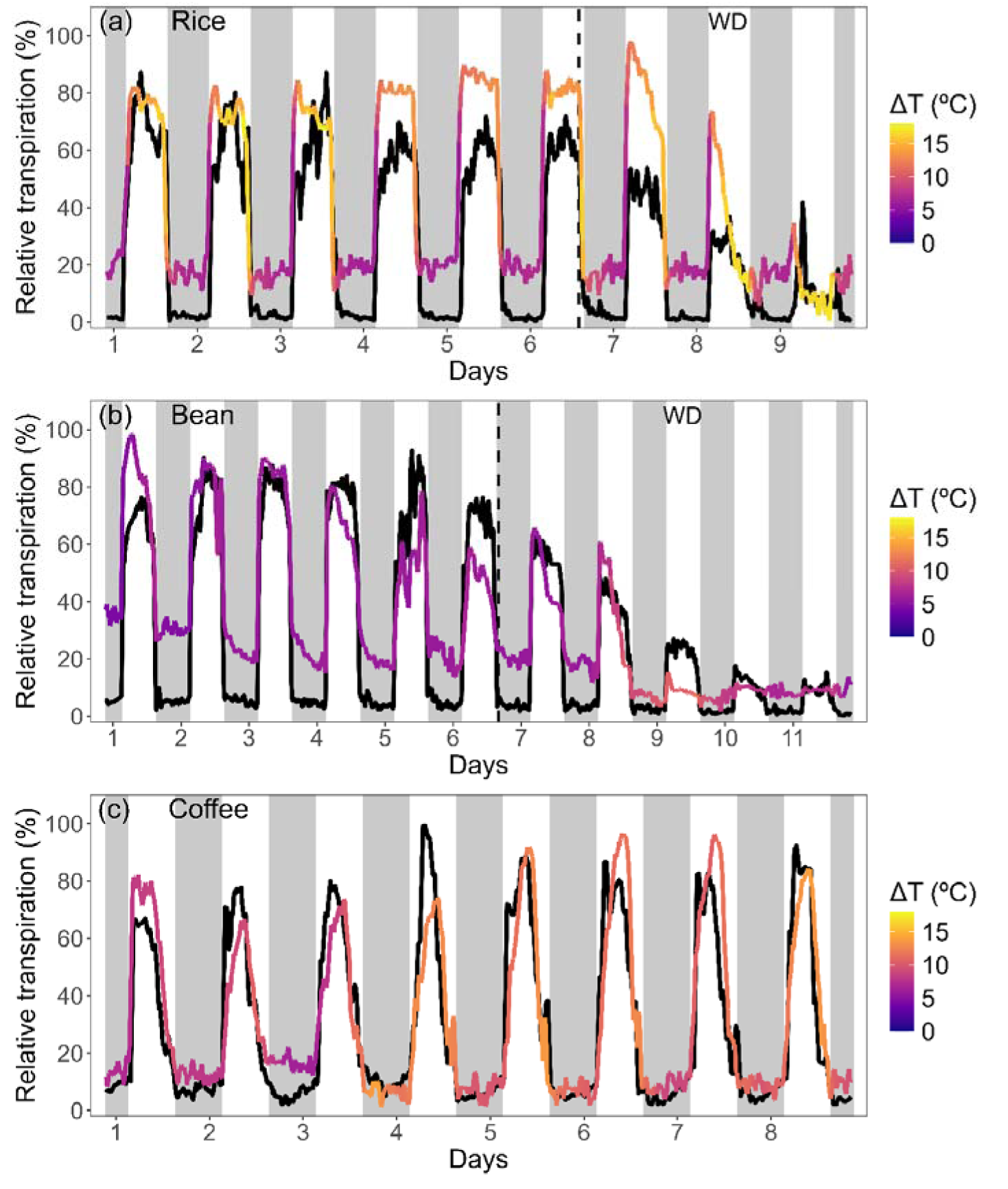
Relative leaf transpiration monitored with the IRGA system (black lines) and the FylloClip (coloured lines) for rice (a), common bean (b) and coffee (c) plants under artificial light but variable air temperature and relative humidity. Measurements were taken every 15 minutes on the same leaf of a single plant. The FylloClip line is coloured according to the dew point depression value (ΔT), with higher values shown in yellow and lower values in blue. For rice and common bean plants, watering was stopped on day 6, causing a period of water deficit (WD; vertical dashed line). Grey background areas represent dark periods (night-time), while white areas represent periods when the light was on (daytime).

When irrigation was suspended for rice and common bean plants, both IRGA and FylloClip values were reduced, with transpiration reaching values close to zero after 3 days of water withholding for rice, and five days for common bean plants (Fig. 3a,b). For rice and common bean, a slight divergence in the night-time baseline was observed between the relative measurements obtained with the FylloClips and those obtained with the IRGA, particularly under low ΔT. This pattern was not observed for coffee, which also exhibited consistently higher ΔT values throughout the experiment. Additionally, higher values of transpiration were recorded for rice with the FylloClip compared to the IRGA between days 4 and 8.

We found a linear correlation between the FylloClip and IRGA data when considering the full dataset across all three species (Pearson’s *r* = 0.85; Fig. 4a). When this analysis was restricted to daytime data, the correlation remained significant (*r* = 0.76; Fig. 4b). Coffee showed the closest agreement between both approaches, with values consistently aligned along the 1:1 line (Fig. 4). Common bean showed a slight deviation from this line at night in the full dataset when IRGA-derived transpiration approached zero (Fig. 4a), consistent with the baseline differences described above (Fig. 3b). In rice, a similar pattern is observed during night-time and also extends to some daytime measurements (Fig. 4b), when the FylloClip values tend to be higher than those obtained with the IRGA.

**Figure 4:**
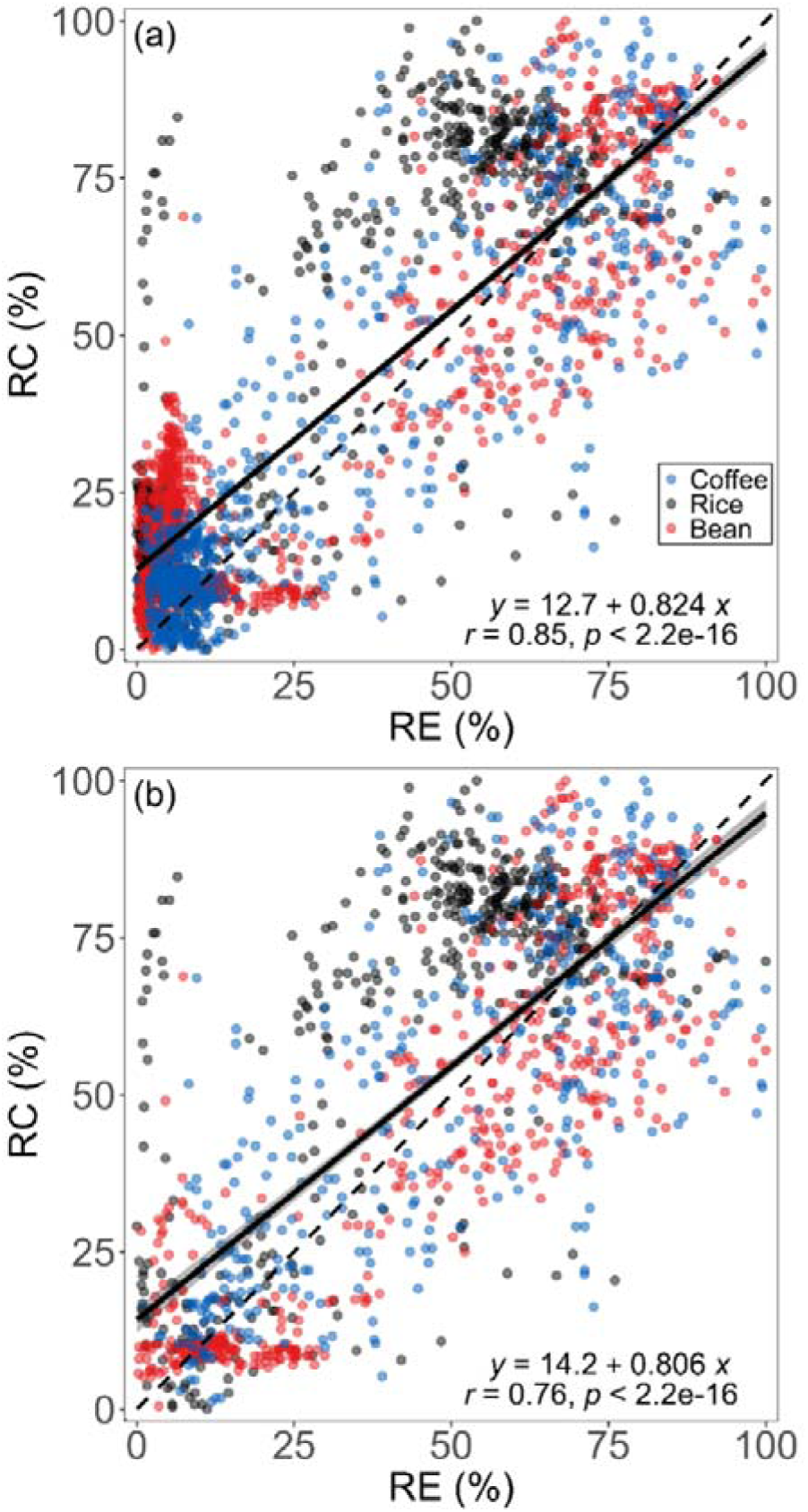
Relationship between relative capacitance measured with Fylloclips (RC) and relative transpiration (RE) measured with an IRGA system, with all day and night-time measurements included (a), or daytime measurements only (b). Each point represents paired measurements of the relative capacitance (RC) and relative transpiration (RE) obtained simultaneously on a single leaf of coffee (*C. arabica*, blue), rice (*O. sativa*, black) and common bean (*P. vulgaris*, red) (*n* = 2,581 measurements in “a” and *n* = 1,373 measurements in “b”). The solid line represents the linear fit, with the shaded area indicating the confidence interval, while the dashed line denotes the 1:1 relationship. The fit equations, Pearson’s *r*, and *p*-values are shown in the graph.

FylloClips were installed on eight coffee plants under greenhouse conditions in experiment 3. Inside the greenhouse, the temperature ranged from 19 to 33.6 °C and RH from 42.7 to 100% (Fig. S3a). During the night, RH inside the greenhouse frequently reached 100%, with both ΔT and VPD often approaching zero (Figs. 5 and S3). The maximum ΔT recorded during the experiment was 14.4 °C (Fig. 5) and the maximum VPD was 3 kPa (Fig. S3b). The relative capacitance measured with the FylloClip showed a daily pattern, increasing in the morning after sunrise and decreasing after sunset. Unlike what was observed in the experiments 1 and 2, on most nights, a slight increase in RC occurred toward the end of the night, during the early morning hours, when ΔT decreased and approached zero (Fig. 5). In contrast, on nights of days 10 and 11, this increase began earlier, already in the late afternoon (Fig. 5), as RH was close to 100% (Fig. S3a).

**Figure 5:**
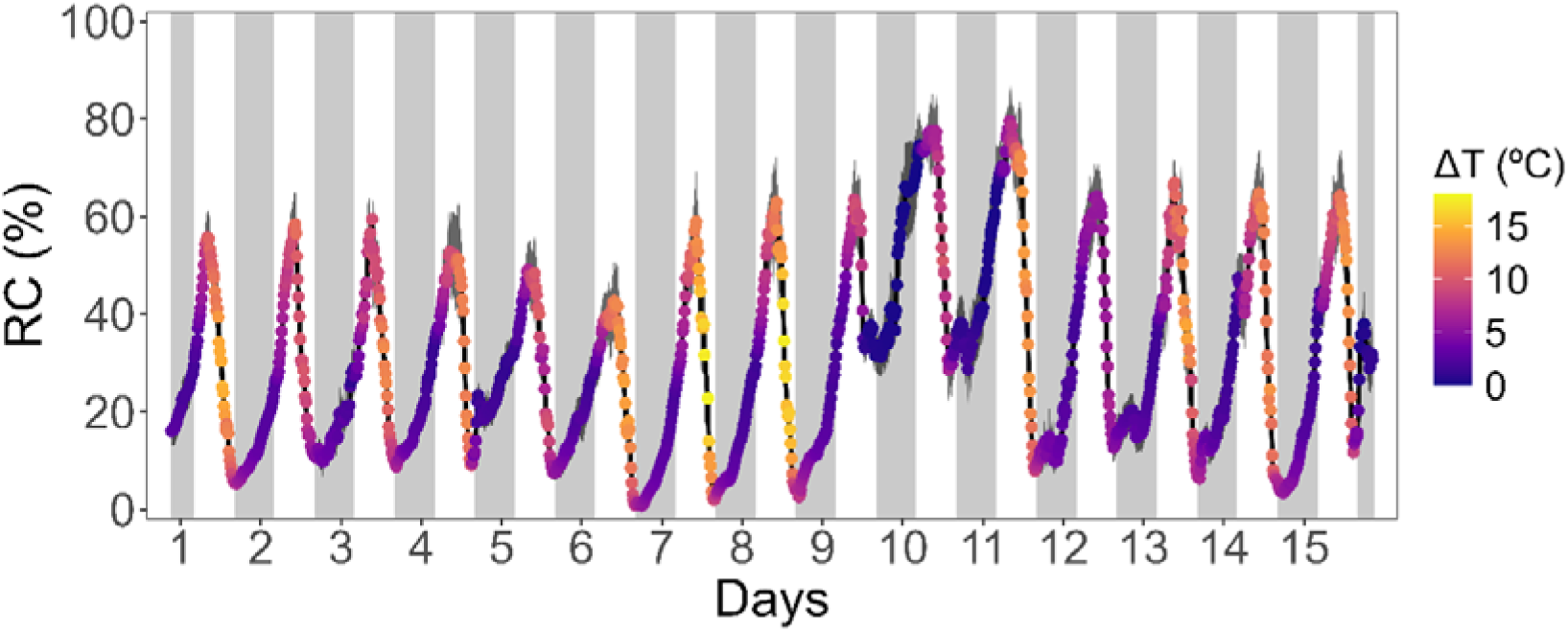
Relative capacitance monitored with FylloClips every 15 minutes in coffee (*C. arabica*) plants under greenhouse conditions for 15 days, with natural variation of light and air temperature and relative humidity. Each symbol represents mean values (*n* = 8 plants, with one sensor per plant), and the dark grey shadow represents the standard error. Symbols are coloured according to the dew point depression value (ΔT), with higher values shown in yellow and lower values in blue. Grey background areas represent night-time, while white areas represent daytime.

FylloClips were also installed on coffee plants under field conditions for one month (experiment 4). Over the course of the experiment, the air temperature ranged from 17.8 to 36.5 °C, and RH from 34 to 100% (Fig. S4a). Similar to the greenhouse conditions, RH frequently reached 100% during the night in the field, with both ΔT and VPD often approaching zero (Figs. 5 and S4). Maximum ΔT values reached 18 °C (Fig. 5) and VPD reached 4 kPa (Fig. S4b). Rainfall occurred on most days after the first week of the experiment, including a heavy rainfall event on the 14^th^ day (Fig. 6b). When heavy rainfall occurred, the FylloClip sensors showed increased values of RC at night (Fig. 6). However, elevated RC values were not restricted to rainfall events. For instance, 2 mm of rainfall occurred in the afternoon on day 16 (Fig. 6b); although no rainfall was recorded during the night, RC values remained high, as RH exceeded 95% (Fig. S4) and ΔT remained close to zero. Similarly, no rainfall was recorded on days 20, 21, and 22, yet RC increased during the night under low ΔT conditions (Fig. 6a). Moreover, heavy rainfall did not cause the FylloClips to detach from the leaves.

**Figure 6:**
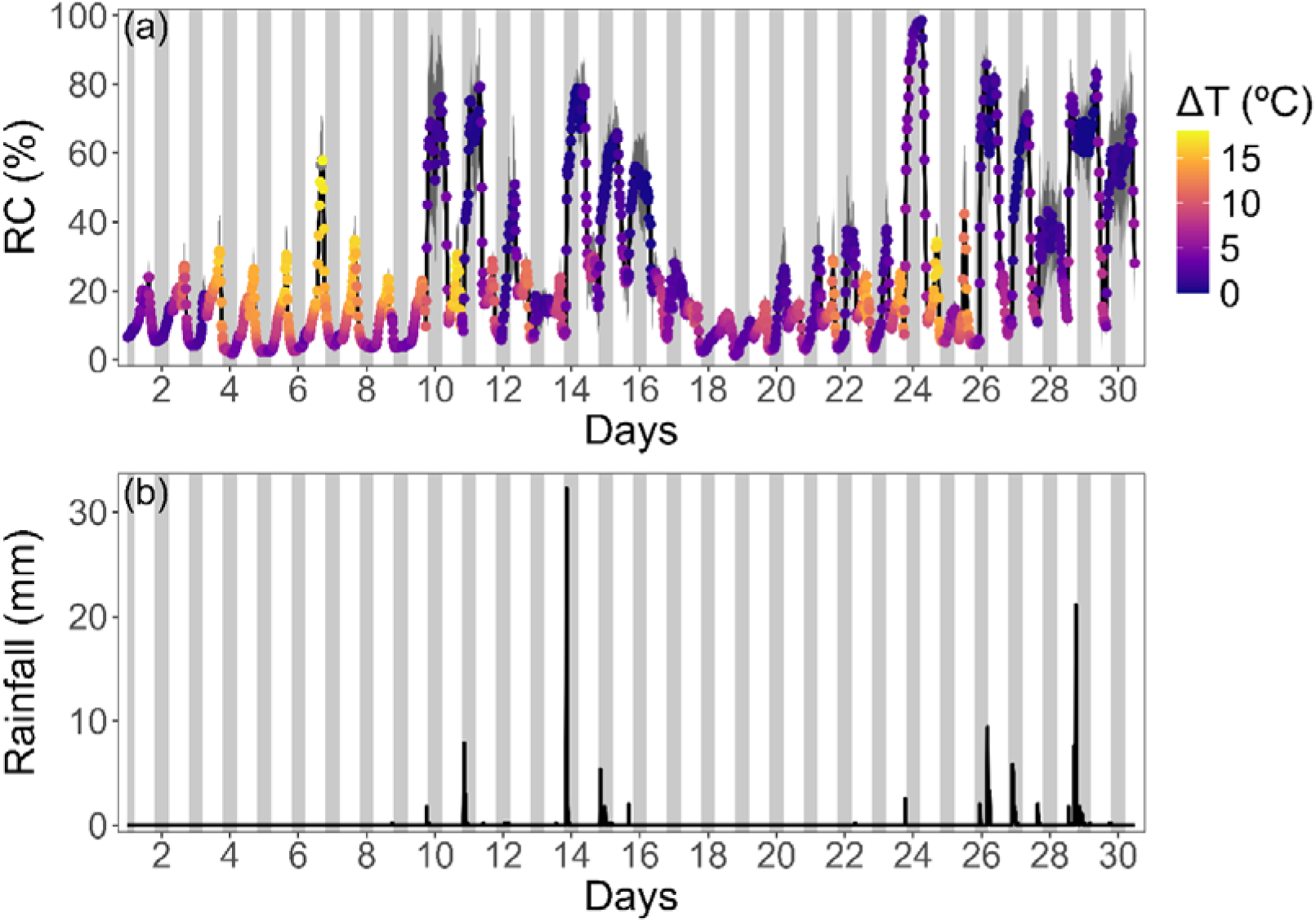
Relative capacitance (RC, %) monitored with FylloClips on coffee plants under field conditions (a); and rainfall recorded at the experimental site (b). Each symbol represents a mean value based on four plants (with one sensor per plant), and the dark grey shaded area indicates the standard error. Symbols are coloured according to the dew point depression value (ΔT), with higher values shown in yellow and lower values in blue. In (b), bars represent rainfall in mm measured every 15 minutes. Grey background areas indicate night-time periods, whereas white areas indicate daytime periods.

Based on a segmented regression of the data from experiments 3 and 4, we identified a breakpoint in the relationship between relative capacitance and ΔT measured during the night, indicating a change in the slope of the relationship (Fig. 7). Below this breakpoint, RC decreased sharply with increasing ΔT, whereas the decline became more gradual above the breaking point. For coffee plants under greenhouse conditions (Experiment 3), the breakpoint occurred at ΔT = 2.80 ± 0.06 °C (Fig. 7a), while under field conditions (Experiment 4) it was 2.97 ± 0.10 °C (Fig. 7b). Since transpiration at night was assumed to be negligible, this breakpoint was used to determine the ΔT range over which RC was largely influenced by condensation on the sensor, and not related to transpiration. Visual observation did not show any evidence of guttation at the leaf margins or leaf tips in coffee plants.

**Figure 7:**
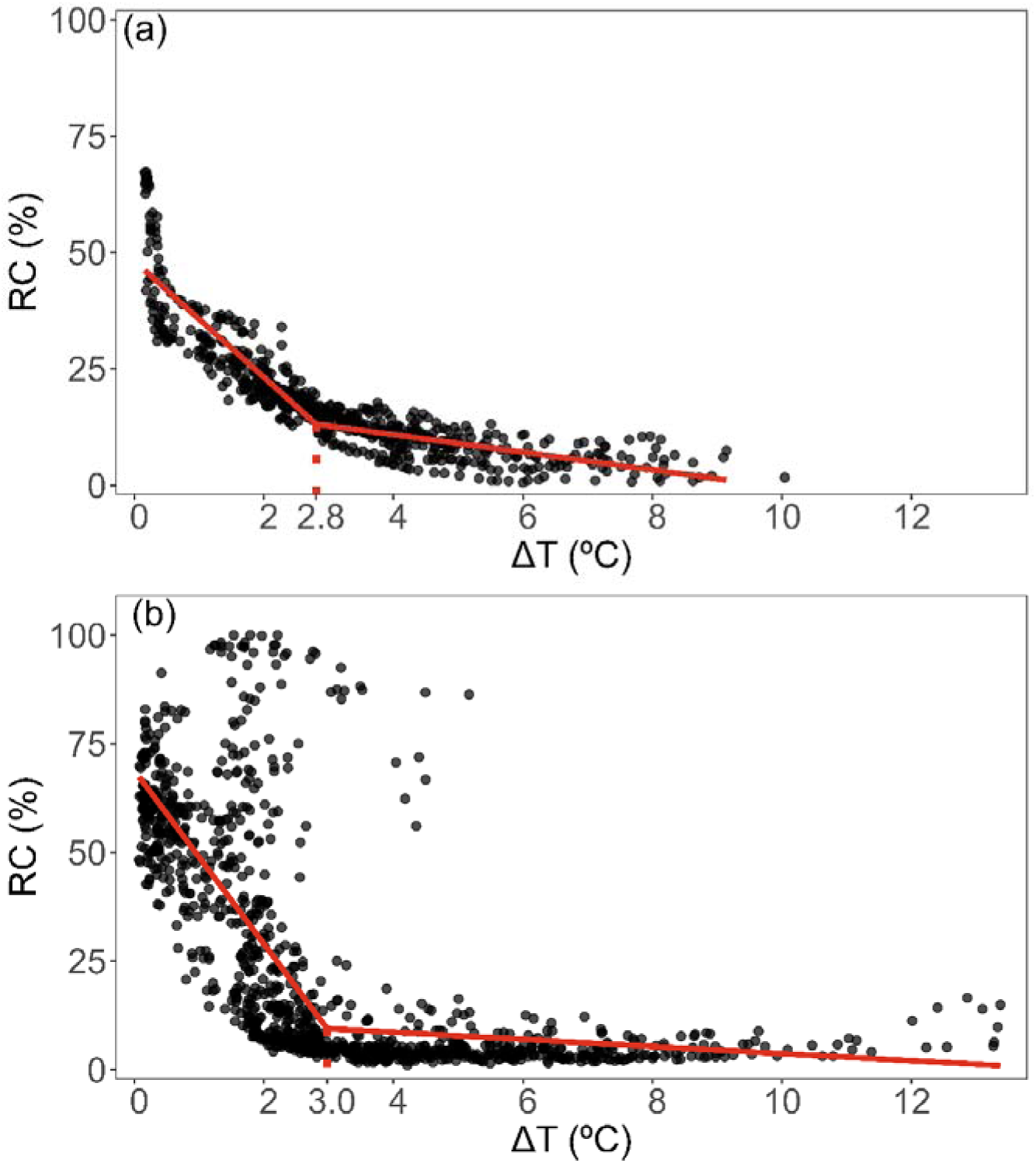
Relationship between relative capacitance (RC) and the difference between air temperature and dew point temperature (ΔT) for coffee (*C. arabica*) plants under greenhouse (a) and field (b) conditions. Black symbols represent mean night-time observations (averaged across sensors), with *n* = 690 in (a) and *n* = 1140 in (b). The means were based on a total of eight and four plants for greenhouse and field conditions, respectively. The red line indicates the fitted segmented regression, showing a breakpoint at ΔT of 2.8 °C in (a) and 3.0 °C in (b), where the slope of the relationship changes.

## Discussion

### FylloClips capture transpiration dynamics at high temporal resolution

Our results provide clear and convincing evidence that the FylloClip sensor captures the daily dynamics of leaf transpiration across all experiments. To our knowledge, this is the first time that a direct comparison between measurements taken with a FylloClip and an infra-red gas analyser has been made. Intriguingly, there was a close agreement between the FylloClip and IRGA-based measurements of leaf transpiration, which demonstrates that the FylloClip data represent a valid and useful proxy for estimating transpiration (Fig. 3; Thalheimer 2023; Thalheimer *et al*., 2026). Therefore, the validation of this sensor opens up a wide range of possibilities to investigate transpiration and stomatal dynamics of plants in an unprecedented, automated way, and at a high temporal resolution over long time periods. Moreover, achieving large-scale measurements at a high spatial resolution with many sensors installed simultaneously should also be feasible given the low cost of the sensor and its potential to use under a wide range of environmental conditions, including high temperature, low VPD, and low ΔT.

The sensor capacitance increased shortly after lights were turned on and decreased when these were turned off, suggesting a close association between sensor capacitance, stomatal behaviour, and leaf gas exchange (Fig. 2; McAdam *et al*., 2016). All eight sensors displayed consistent patterns, and similar responses were also observed in plants of rice (*O. sativa*), common bean (*P. vulgaris*), and coffee (*C. arabica*) under variable environmental conditions (Fig. 3). When irrigation was suspended, a gradual reduction in transpiration was observed both in IRGA- and FylloClip-measurements (Fig. 3a,b). However, the FylloClip sensor appeared to be slightly less sensitive than the IRGA device when transpiration rates became very low in common bean plants under water deficit. Under these conditions, the FylloClip readings approached zero, while the IRGA still detected very low transpiration rates (Fig. 3b). Both methods also captured distinct diurnal patterns among species. In common bean and rice, transpiration increased after the onset of light and remained relatively high throughout the day. In contrast, coffee exhibited a clear midday maximum followed by a decline in transpiration during the afternoon, even while light conditions remained constant.

We observed a shift in the baseline values of the FylloClips, which were recorded during night-time when transpiration is very low to absent, especially between the beginning and the end of the experiments with rice, common bean and coffee plants (Fig 3). These changes can largely be explained by daily differences in environmental conditions. At the beginning of the experiment, night-time ΔT values were lower than those observed later (Fig. 3) due to higher RH and temperature (Fig. S2). Such high night-time capacitance values may result from incomplete evaporation of water accumulated on the sensor surface during daytime transpiration. Because the sensor is positioned close to the leaf surface (depending on the leaf thickness, leaf micromorphology, and the clamping force the sensor clip), the restricted air exchange within the narrow gap between the leaf and the sensor plate may slow the drying process, allowing residual moisture to persist overnight. When ΔT was high, the baseline values of the FylloClip were low, which started around the third day for common bean, and on the eighth day for coffee plants. These shifts in the baseline values of the FylloClip were likely caused by drier air conditions (Fig. S2), which enhanced evaporation and facilitated the removal of moisture previously accumulated in the gap between the sensor plate and the leaf surface.

In addition to the baseline shifts, species-specific differences became evident when comparing FylloClip and IRGA measurements. Coffee showed the strongest agreement between approaches, with values closely following a 1:1 relationship across conditions (Figs. 3c and 4) and indicating that the FylloClip reliably captured relative changes in transpiration under the tested conditions. In contrast, rice exhibited a consistent tendency for higher transpiration estimates derived from the FylloClip compared to the IRGA, both under low-transpiration conditions and during the daytime (Figs. 3a and 4). Rather than reflecting true differences in transpiration rates, this apparent overestimation is more likely associated with differences in leaf surface properties and boundary layer characteristics that influence condensation dynamics at the sensor interface. Some mechanisms may explain these species-specific patterns. First, differences in leaf surface properties, such as wettability and cuticle structure, may affect water retention and evaporation near the sensor. In addition, the presence of small trichomes on leaves or other leaf surface structures may act as nucleation sites for microdroplet formation under high humidity (Roth-Nebelsick *et al*., 2012; Burkhardt & Hunsche, 2013; Konrad *et al*., 2015). Finally, differences in leaf–sensor contact among species may influence the extent to which water accumulates or persists within the measurement gap. Together, these factors could enhance local condensation independently of actual transpiration, leading to higher apparent signals from FylloClip. These findings highlight that, while the FylloClip effectively tracks relative changes in transpiration, species-specific leaf traits and microenvironmental conditions at the sensor interface must be considered. Further work exploring a broader range of species and leaf morphologies would help disentangle these effects and refine the interpretation of FylloClip measurements, including the potential effect of nocturnal transpiration and/or guttation (Marks & Lechowicz, 2007; Singh, 2016).

Transpiration rate is not only driven by stomatal behaviour, but also to some extent by the resistance of the boundary layer. The thin layer of air surrounding a leaf surface can limit air movement considerably, and reduce convective exchange between stomata and the environment (Defraeye *et al*., 2013). Because the FylloClip is directly in contact with the leaf, a tight gap between the sensor plate and the leaf may create a semi-enclosed air space, which may reduce ventilation and favour local saturation and condensation. Under such conditions and particularly when ΔT and VPD are low, water vapour may accumulate near the leaf surface, promoting local saturation and condensation. This mechanism likely explains why transpiration rates estimated by the FylloClip can be higher than those measured by the IRGA, especially under low evaporative demand (Fig. 3). The onset of condensation on a sensor that is tightly connected to a leaf could also be promoted by the presence of small ridges, pores, or surface irregularities of the sensor and/or printed circuit board. It is known that the Kelvin effect, which describes how the local vapour pressure above a liquid is affected by its curvature, may contribute to reverse transpiration and even leaf water uptake, although the latter is also affected by various other processes (Vesala *et al*., 2017; Berry *et al*., 2019; Schreel & Steppe, 2020; Matos *et al*., 2024). Very small gaps between the sensor head and the leaf may also act like capillary spaces, retaining thin water films, and reducing evaporation. Therefore, under low ΔT and low VPD conditions, this restricted air exchange may favour the accumulation of water vapour near the leaf surface, increasing the likelihood of condensation. Preliminary tests with a different sensor design and clamping force of the clip suggest that the gap between the sensor and the leaf may influence sensitivity, potentially affecting the sensor response and its susceptibility to condensation (unpublished data). It is possible that very tight contact and a high clamping force of the clip may impose mechanical stress on the leaf tissue, potentially leading to slight damage. Clearly, more research is needed to explore the effect of the sensor head on its measurements. Ideally, the surface properties of the sensor should mimic those of the leaf surface to detect leaf-liquid-vapour-sensor interactions as natural as possible, without side effects.

### Environmental effects on capacitance measurements

When the FylloClip sensors were installed under conditions of variable RH (or VPD) and temperature, daily patterns of relative capacitance could be detected. However, higher environmental variability, and especially high RH, requires sufficient care to interpret the capacitance measurements. We found that the difference between air temperature and dew point temperature (ΔT) was an important environmental factor for sensor performance. When ΔT and VPD approach zero, water vapour can condense on the sensor surface. This is relevant particularly during the night, when air temperature typically decreases while RH increases, bringing air temperature closer to the dew point temperature, therefore reducing ΔT. As discussed by Thalheimer *et al*. (2026), an increase in sensor capacitance occasionally observed at night can also be attributed to dew formation when the ambient temperature reaches the dew point. This effect may especially be relevant in very humid environments, where ΔT frequently approaches zero. Under these conditions, condensation on the sensor plate can increase capacitance values, generating signals unrelated to leaf transpiration. Consistent with this interpretation, our breakpoint analysis indicated that capacitance responses shifted at ΔT values of approximately 2.8 °C in the greenhouse experiment and 3.0 °C under field conditions (Fig. 7), which corresponded to VPD values close to zero. These relatively similar thresholds, despite contrasting environmental settings, suggest that a narrow range of low ΔT and VPD marks the transition at which condensation effects begin to influence sensor readings. In addition to condensation, the formation of guttation droplets via marginal or laminar hydathodes (Schenk *et al*., 2021; Michaud *et al*., 2024) could also result in high FylloClip records at night or early morning, especially in Poaceae species such as rice and maize, although this has not been observed in our experiments.

As could be expected, rainfall is an important environmental factor affecting the FylloClip capacitance measurements under field conditions. During periods of rainfall, increases in relative capacitance were observed. This effect is likely caused by rainwater entering the gap between the sensor plate and the leaf surface. Water accumulating in this space increases the measured capacitance due to the high dielectric constant of water (Thalheimer *et al*., 2026). Consequently, rainfall can generate signals that should not be misinterpreted as transpiration. Furthermore, the FylloClip measurements are strongly affected during periods of frequent rainfall and low ΔT, as observed on the last days in experiment 4 (Fig. 6).

### The FylloClip as a tool for monitoring plant water status

The FylloClip represents a promising tool for monitoring plant water status over time. This sensor fits within the emerging class of plant wearable sensors, which enable continuous and *in situ* monitoring of physiological processes directly at the organ level (Li *et al*., 2024). By positioning sensors directly on plant organs, these technologies allow real-time tracking of physiological responses related to plant performance and health status, providing new opportunities for plant monitoring and phenotyping (Kim *et al*., 2019; Zhang *et al*., 2023). Such monitoring approaches are particularly relevant for irrigation management in precision agriculture. Irrigation scheduling is a widely used approach to manage crop water requirements throughout the growing season. Traditionally, irrigation decisions are based on environmental and soil-related variables, such as evapotranspiration estimates, precipitation, and soil moisture measurements (Dong *et al*., 2024). While these approaches provide valuable information about the soil water availability or the atmospheric demand, they do not necessarily reflect the actual plant water status. For this reason, monitoring leaf transpiration dynamics may represent a promising strategy for irrigation management, as it allows water supply to be adjusted according to the plant’s real-time needs. Monitoring the effectiveness of irrigation systems both in spatial and temporal scales is an example of how FylloClips could provide a useful tool for improving crop yield, increasing water use efficiency in plants both in greenhouses and field conditions.

Although the sensor does not provide quantitative measurements of transpiration directly, its ability to capture consistent diurnal patterns related to stomatal behaviour – the main driver of leaf transpiration – allows the identification of changes in plant water status at a high temporal unprecedented resolution.

## Conclusion

The FylloClip sensor is shown to be a highly promising tool for long-term monitoring of transpiration and dynamics of stomatal behaviour. Its very low cost, operational simplicity, and ability to automatically record data over extended periods make it possible to apply many sensors simultaneously and to capture continuous and large-scale patterns that are related to the water status of plants. By tracking daily dynamics associated with transpiration, the sensor can provide valuable, long-term insights into how plants respond hydraulically to their environment. These characteristics make the FylloClip a highly useful tool for studies requiring high temporal resolution measurements and large sensor replication, such as sensor-based forest monitoring systems. In addition, its easy installation suggests considerable potential for applications in precision agriculture, where continuous monitoring of plant water responses to the surrounding environment could support more informed irrigation management and improve the water use efficiency of plants.

## Acknowledgments

The authors acknowledge the National Council for Scientific and Technological Development (CNPq, Brazil; Grants #304295/2022-1, #446107/2024-7 and #314858/2025-3), the São Paulo Research Foundation (FAPESP, Brazil; Grant #22/04006-2), the Brazilian Federal Agency for Support and Evaluation of Graduate Education (CAPES, Brazil, Grant #88887.078769/2024-00), the Deutsche Forschungsgemeinschaft (DFG, German Research Foundation, project #508216003 and #457287575) and the Office for Gender Equality at Ulm University for support received. LP and MTM thank M. Thalheimer for technical advice with the construction of FylloClips, and SJ acknowledges HJ Schenk for drawing attention to the FylloClip.

## Authors contribution

MTM, LP, SJ, and RVR developed the idea and planned the experiments, which were conducted by MTM, LP, SK, JO, AFDF and CCC. MTM and LP analysed the data. All authors contributed to the discussion and manuscript writing, with substantial inputs from MTM, SJ and RVR.

## Supporting information

**Figure S1:** Environmental conditions of the growth chamber during Experiment 1.

**Figure S2:** Environmental conditions inside a room used for Experiment 2.

**Figure S3:** Environmental conditions inside a greenhouse used for Experiment 3.

**Figure S4:** Environmental conditions in our experimental field during Experiment 4.

## Data availability

The data that support the findings of this study are available within the article and its supplementary material. Raw data and R scripts are available from the corresponding author upon request.

