## Supplementary material for "Continuous monitoring of plant transpiration dynamics with a leaf-mounted sensor across environmental conditions"

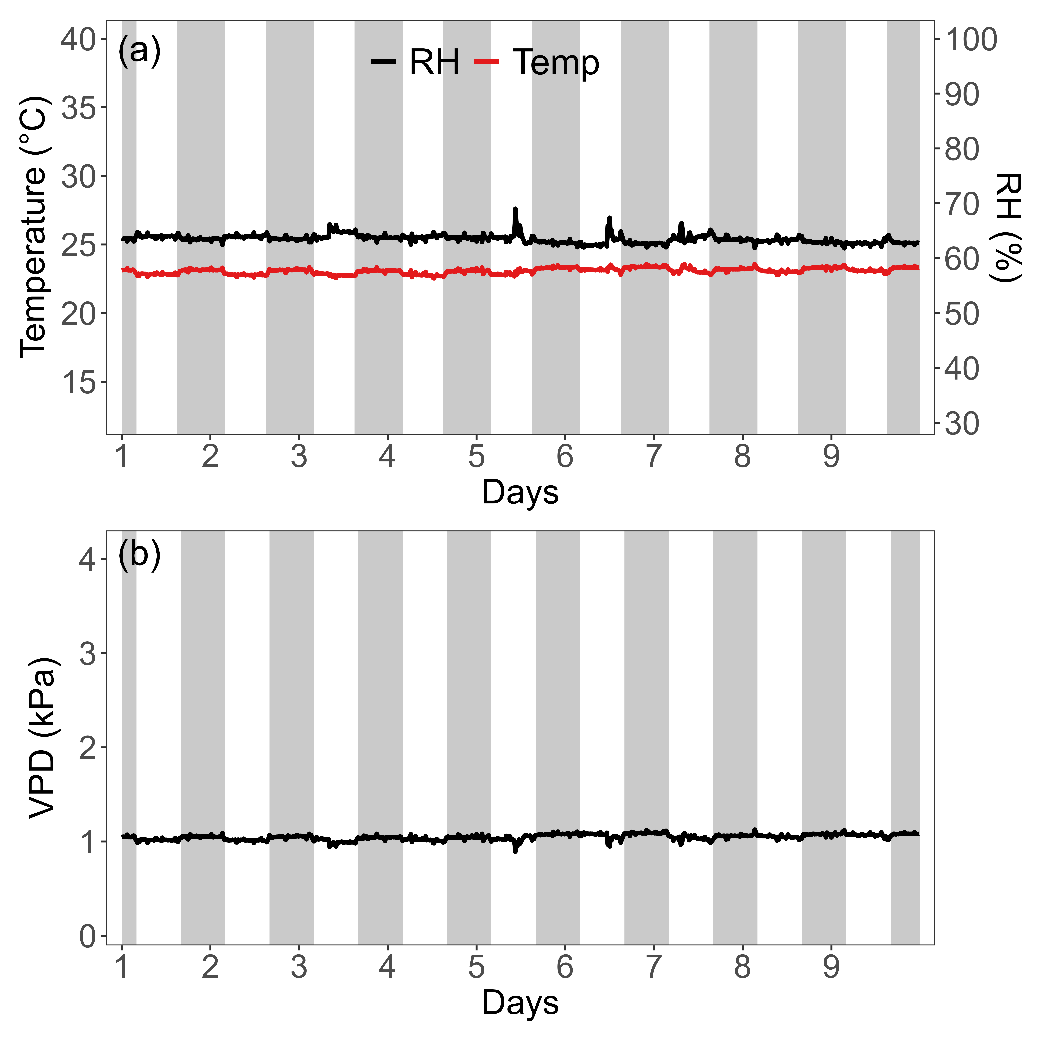

**Figure S1:** Environmental conditions inside the growth chamber during the Experiment 1, with maize plants: air temperature (in red) and relative humidity (RH, in blue) shown in (a); and vapour pressure deficit (VPD) in (b). Grey background areas represent dark periods (night), while white areas represent periods when the light was on (day).

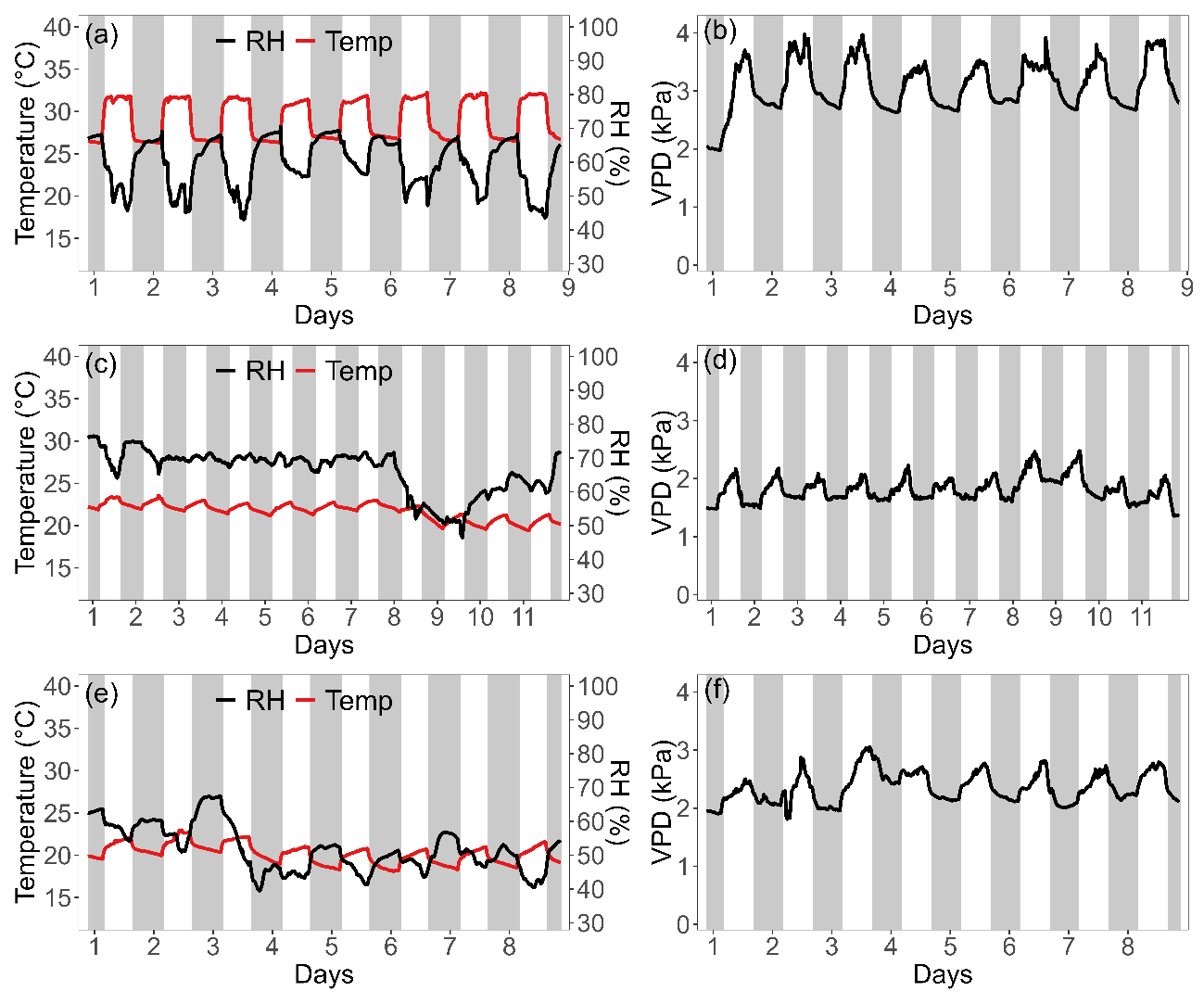

**Figure S2:** Environmental conditions inside a room used for the Experiment 2, with air temperature (in red) and relative humidity (RH, in black) when rice (a), common bean (c), and coffee (e) plants were evaluated. The corresponding air vapour pressure deficit (VPD) is shown in (b), (d), and (f). The experiments were run separately for each species. Grey background areas represent dark periods (night), while white areas represent periods when the light was on (day).

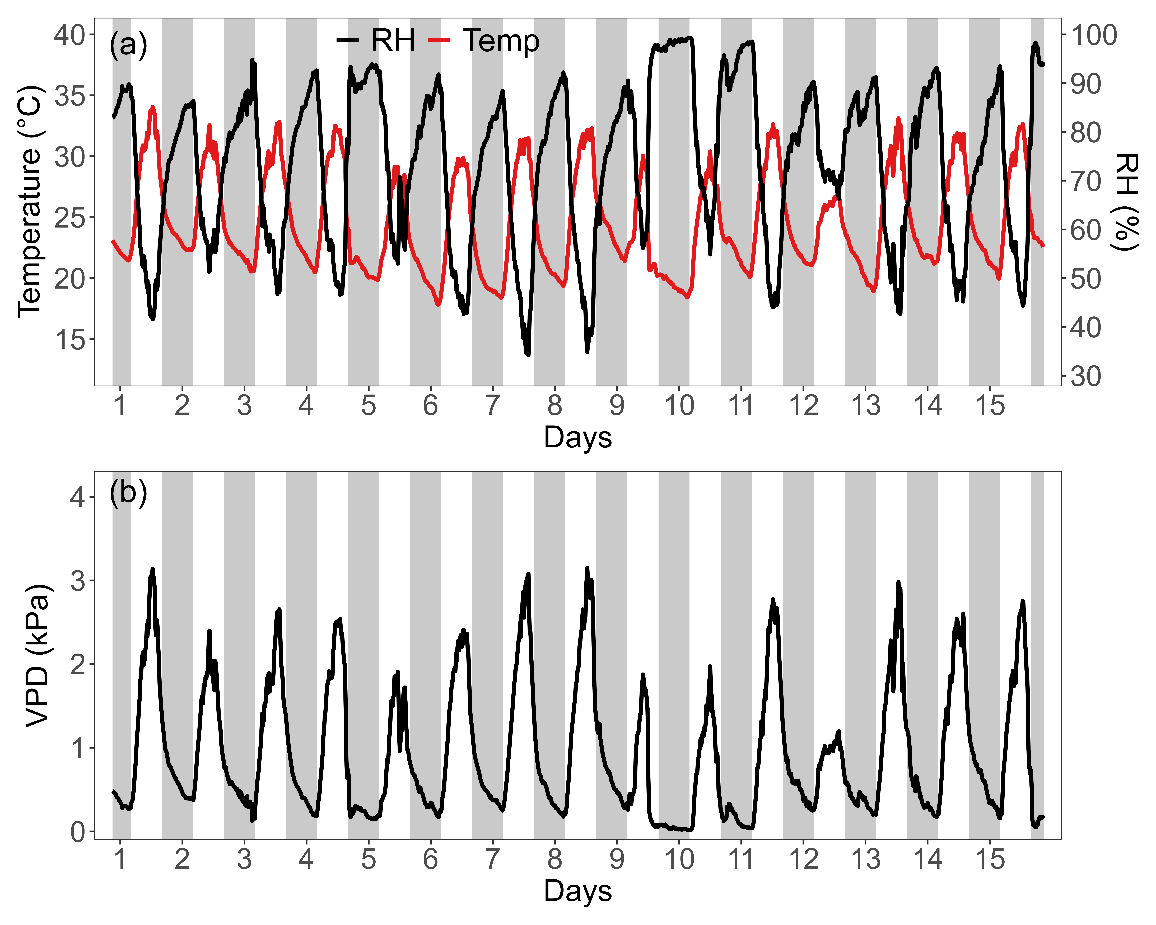

**Figure S3:** Environmental conditions inside a greenhouse used for the Experiment 3, with coffee plants: air temperature (in red) and relative humidity (RH, in blue) in (a); and air vapour pressure deficit (VPD) in (b). Grey background areas represent night periods, while white areas represent day periods.

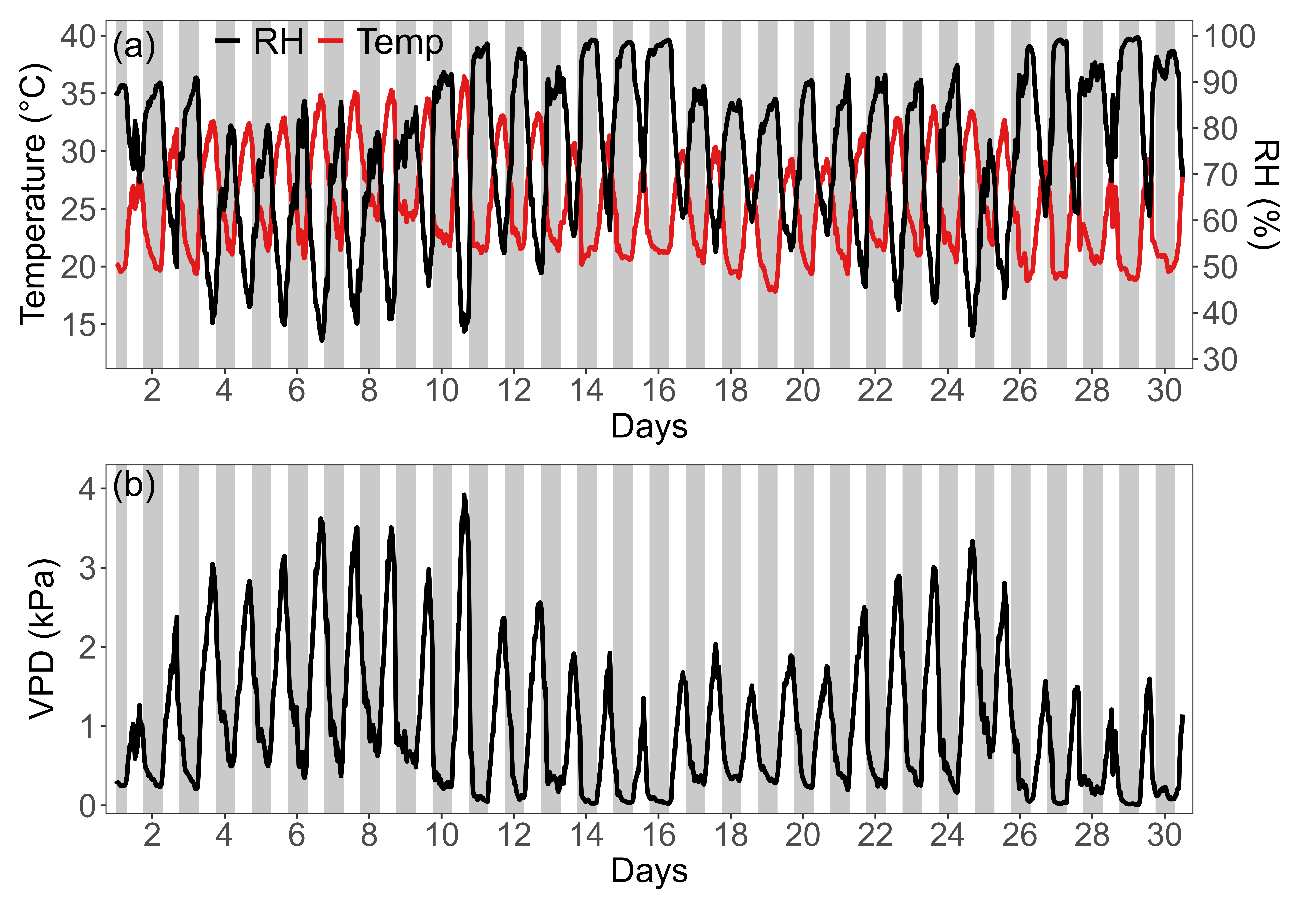

**Figure S4:** Environmental conditions in our experimental field during the Experiment 4, with coffee plants: air temperature (Temp, in red) and relative humidity (RH, in blue) in (a); and air vapour pressure deficit (VPD) in (b). Grey background areas represent night periods, while white areas represent day periods.
